# Utilizing Pre-trained Network Medicine Models for Generating Biomarkers, Targets, Re-purposing Drugs, and Personalized Therapeutic Regimes: COVID-19 Applications

**DOI:** 10.1101/2023.02.21.527754

**Authors:** Jianghui Xiong

## Abstract

In this paper, we present a novel pre-trained network medicine model called Selective Remodeling of Protein Networks by Chemicals (SEMO). We divide the global human protein-protein interaction (PPI) network into smaller sub-networks, and quantify the potential effects of chemicals by statistically comparing their target and non-target gene sets. By combining 9607 PPI gene sets with 2658 chemicals, we created a pre-trained pool of SEMOs, which we then used to identify SEMOs related to Covid-19 severity using DNA methylation profiling data from two clinical cohorts. The nutraceutical-derived SEMO features provided an effective model for predicting Covid-19 severity, with an AUC score of 81% in the training data and 80% in the independent validation data. Our findings suggest that Vitamin D3, Lipoic Acid, Citrulline, and Niacin, along with their associated protein networks, particularly STAT1, MMP2, CD8A, and CXCL8 as hub nodes,could be used to effectively predict Covid-19 severity. Furthermore, the severity-associated SEMOs were found to be significantly correlated with CD4+ and monocyte cell proportions. These insights can be used to generate personalized nutraceutical regimes by ranking the relative severity risk associated with each SEMO. Thus, our pre-trained SEMO model can serve as a fundamental knowledge map when coupled with DNA methylation measurements, allowing us to simultaneously generate biomarkers, targets, re-purposing drugs, and nutraceutical interventions.

## Introduction

The COVID-19 pandemic caused by the severe acute respiratory syndrome coronavirus 2 (SARS-CoV-2) has posed a tremendous challenge to existing medical research, drug development and clinical nursing. Variations in symptom presentation between individuals suggest complexity in virus-host interaction. Post-acute sequelae of COVID-19, referred to as ‘long COVID’, have been identified with over 200 symptoms impacting multiple organ systems [1]. Traditionally, diseases have been defined largely by their phenotypes. However, more novel approaches that capture intrinsic and casual mechanisms could revolutionize the way we define, diagnose, and treat diseases [2].

Current signaling pathways are mostly hand-curated by human experts. Thus, a data-driven approach to generate disease pathology and drug mechanism of action would be invaluable for emerging diseases. To this end, large-scale omics studies of clinical cohorts can provide comprehensive molecular characterizations of COVID-19 through epigenomics, transcriptomics, proteomics, genomics, lipidomics, immunomics and metabolomics [3, 4]. These data are a rich source for the development of disease biomarkers. Repurposing methods combining disease profiling data and compound features have shown promise in quickly prioritizing candidate drugs for validation[5].

Considering immune dysregulation is a primary pathology of COVID-19 and its sequelae, we hypothesize that a health status that quantifies both the state of immune centric protein interaction networks and their potential to be regulated by chemicals would facilitate the generation of a disease knowledge map in a data-driven way. To this end, the novel network module SEMO (selective remodeling of protein networks by chemicals) was proposed. Module activity, referred to as the SEMO index, can be measured by omics methods such as whole blood DNA methylation profiling. Potential regulation by endogenous metabolites, nutrient supplementations, nutraceuticals, approved drugs, or natural products can be encoded in the module’s definition. Whole blood DNA methylation profiling combined with a computational biology pipeline would enable the simultaneous output of three things: (1) disease/severity biomarkers, (2) therapeutic targets, and (3) candidate drugs. Whole blood DNA methylation profiling is preferred to use as the input of generative modeling due to its precision, widespread use in human aging regression (so-called “DNA methylation clock”)[6] and high-resolution immune cell profiling [7].

In this paper, we introduce the concept of SEMO, explore its association with Covid-19 disease severity, and interrogate its dynamic effect on various stages of Covid-19. Using machine learning algorithms, we construct a severity prediction model with SEMO as input predictors and validate it with independent cohorts. We further calculate personalized nutraceutical intervention regimes based on personal DNA methylation profiling derived SEMO profiling. Additionally, we extend the method to FDA-approved drugs and herb chemicals often used in traditional Chinese medicine, and develop a drug repurposing tool based on SEMO calculation. In conclusion, we explore if the combination of DNA methylation profiling, SEMO calculation and machine learning can simultaneously output biomarkers, drug targets, and candidate drugs or interventions for complex diseases.

## Methods

### DNA methylation profiling data of COVID-19

For this study, we analyzed DNA methylation profiles relating to Covid-19 disease severity across two clinical cohorts. Data 1 comprised 473 Covid-19 positive and 101 negative individuals whose bisulphite converted DNA samples were hybridized to the Illumina Infinium MethylationEPIC Beadchip (850k). This data can be obtained freely from the GEO database using the accession GSE179325 [8]. Individuals were classified based on the WHO clinical ordinal scale: PCR negative individuals (uninfected, 0 scale), mild PCR positive individuals (ambulatory or hospitalized with mild symptoms, 1–4 scales), and severe PCR positive individuals (hospitalized with severe symptoms or died, 5–8 scales).

Data 2 was a genome-wide methylation profiling of 164 Covid-19 positive individuals, 296 Covid-19 negative patients, and 65 other respiratory infection samples profiled on a custom Illumina Infinium Methylation BeadChip array. Only data from probes found on Human MethylationEPIC BeadChip (850k) was used for this analysis, and the data can be accessed on the GEO database under the accession GSE16720225 [9]. COVID-19 disease severity was represented by an ordered severity score of (1) discharged from emergency department (home care); (2) admitted to inpatient care; (3) progressed to ICU; and (4) death.

### Identification of COVID-19 associated protein interaction subnetworks

A collection of protein-protein interactions was integrated from the Human Protein Reference Database (HPRD, www.hprd.org). This whole network was then divided into separate hub gene sets. For example, the gene set “p53.n” represents a gene set that includes all genes that have interactions with the protein p53 in the protein-protein interaction networks. The “size” of a gene set is equal to the number of interacting genes it includes. Here, only hub gene sets with 20 or more interacting genes were used.

We generated a gene set that was related to severity of the data set 1. For each gene, the average of all promoter CpG sites (TSS 1500 and TSS 200) were computed as a representative DNA methylation score. Then, we conducted a two-class t-test to filter out genes that significantly differentiated between the two clinical groups (severe and mild group in data1). The gene list was ranked according to the t-test p-values, and the most differentially-expressed genes were utilized as the severity-related gene set. Using the same procedures, we created a severity-related gene set for Data 2. Patients with severity scores of 2 or higher were categorized in the “severe group”, while those with scores of 1 were labeled as the “mild group”. The overlapping of the severity gene sets from Data 1 and Data 2 were used to form the “common severity gene set”.

For each hub gene set (PPI gene set), an association with disease severity was calculated. For a PPI gene set that includes n genes, if it includes k genes that overlap with the previously identified “common severity gene set”, then the severity score of the PPI gene set is represented by the ratio k/n. The severity scores of all PPI gene sets were calculated, sorted in descending order, and the top 100 PPI gene sets were used in further procedures.

### Chemical-PPI targeting significance

Only chemical-PPI gene set combinations which show significant overlap were used to calculate SEMO index. This chemical-PPI targeting significance was represented by -log10(p), where p is the p-value obtained from the hypergeometric test run on each pair of PPI gene set and chemical target gene set; p is calculated by:

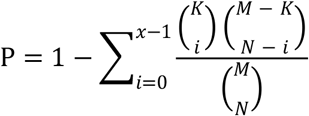

Here, x represents the number of overlapped gene sets in the PPI gene set and given chemical target gene set; K is the number of genes in PPI gene set, N is the number of genes in chemical target gene set, and M is the number of all genes in the genome. Only the PPI-chemical pairs with hypergeometric p-value smaller than 0.01 were used to calculate SEMO index.

### SEMO (Selective remodeling of protein networks) index calculation

Considering PPI gene set ***i***, which has multiple genes, then the intersection of chemical ***j*** target gene set with PPI gene set ***i*** is ***x***, and the different set (belongs to PPI gene set **i** but not in chemical target gene set ***j***) is **y**. The two-class t-test of gene scores (here, DNA methylation score) between set ***x*** with set ***y*** can be calculated by t statistics:

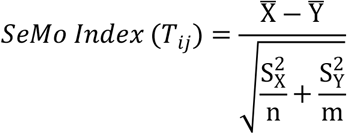

Here, 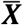 is the mean of gene scores of gene set ***X***, 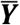 is the mean of gene scores of gene set ***Y***. 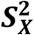 is the variance of gene set X scores, 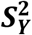 is the variance of gene set ***Y*** scores, ***n*** is the number of genes in set ***X***, and ***m*** is the number of genes in set ***Y***.

Therefore, for any combination of PPI gene set ***i*** with chemical ***j***, the above calculated ***Tij*** is used to represent the state of SEMO (PPI-chemical), for example, “***P53*.*n-Vitamin A***” represents the SEMO related to PPI gene set “P53” and chemical “Vitamin A”.

### Severity associated SEMO calculation

In order to identify severity-associated SEMO features, a combination of three types of chemical targets was scanned: (1) nutraceuticals from DrugBank (https://www.drug-bank.ca), (2) FDA-approved drugs from DrugBank (https://www.drug-bank.ca), and (3) chemicals from traditional Chinese medicine herbs (TCMID, http://www.megabionet.org/tcmid/). The target gene sets for each of these chemicals were downloaded from the STITCH database (http://stitch.embl.de) with a STITCH score of ≥ 200.

All combinations of 100 severity-associated PPIs with the chemical gene sets were then iterated. For the DNA methylation profiling data of each patient in the first dataset (Data 1), SEMO index ***Tij*** were calculated for each PPIs set ***i*** with each chemical gene set ***j***. This resulted in a genes×samples matrix which was transformed into the SEMOs×samples matrix of Data 1. Each SEMO feature was then checked for significant difference between COVID-19 severe and mild groups using a t-test. The SEMO features were then ranked by t-test p-values and formed the list of severity-associated SEMO. The same procedures were conducted on the second dataset (Data 2).

### Severity prediction models

A severity prediction model was developed based on lasso (Least Absolute Shrinkage and Selection Operator) on Data 1 and validated in an independent cohort of 525 patients (Data 2). Three models were generated for the three types of SEMO: those from the combination of PPIs and (1) nutraceuticals, (2) TCM chemicals, and (3) approved drugs. For the nutraceuticals, the intersection of the top 150 severity-associated SEMOs from Data 1 and the top 150 SEMOs from Data 2 was checked. A total of 49 overlapping SEMOs were identified. Based on this 49 SEMOs× samples matrix on Data 1, a classification model was trained for class label 1 (severe group) and class label 0 (mild group) using lasso algorithms. The area under the receiver operating characteristic curve (AUC) were used to quantify the performance of the models.

### Personalized risk profiling and interventions

From a personal perspective, it is essential to analyze all the SEMO features and assess their relative impacts and risks in order to identify the riskiest ones and develop tailored potential interventions and preventative measures. To do this, we used a logistic regression fitting model with a phenotype label of y (severe cases labeled y =1, mild cases labeled y =0) for each severity associated SEMO. Subsequently, each SEMO index was converted into a Risk Score between 0 and 1, resulting in a Risk Score×Patients matrix conversion from the SEMOs×Patients matrix.

### SEMO - immune cell proportion association

To assess the correlation of the severity-predictive SEMO feature with immune cell profiling, an enhanced DNA methylation library data was downloaded from the GEO Database (accession GSE167998) that included libraries to deconvolute peripheral blood [7]. The data used in this study was comprised of 12 different leukocyte subtypes (neutrophils, eosinophils, basophils, monocytes, B cells naïve and memory, CD4+ and CD8+ naïve and memory cells, natural killers, and T regulatory cells), assayed using the Illumina HumanMethylationEPIC array. Twelve artificial mixtures (labeled as “Mix” in the dataset) were hybridized to the Illumina Infinium HumanMethylationEPIC beadchip. The SEMO index was calculated using previously established procedures, and Pearson correlation coefficient was used to measure the correlation between the SEMO index and the cell proportions.

## Results

### Pre-trained SEMO library and the information processing workflow

Here, a protein-protein interaction (PPI) subnetwork was used as the protein network feature. We calculate the statistical difference between the interaction of the PPI gene set and the chemical target genes (x) and the PPI gene set that does not overlap with the chemical target genes (y) (see **Figure 1A**). If the DNA methylation values of x and y show statistically significant difference between severe and mild patients, this SEMO feature is meaningful in terms of discriminating phenotype (disease severity, **Figure 1B**).

**Figure 1.**
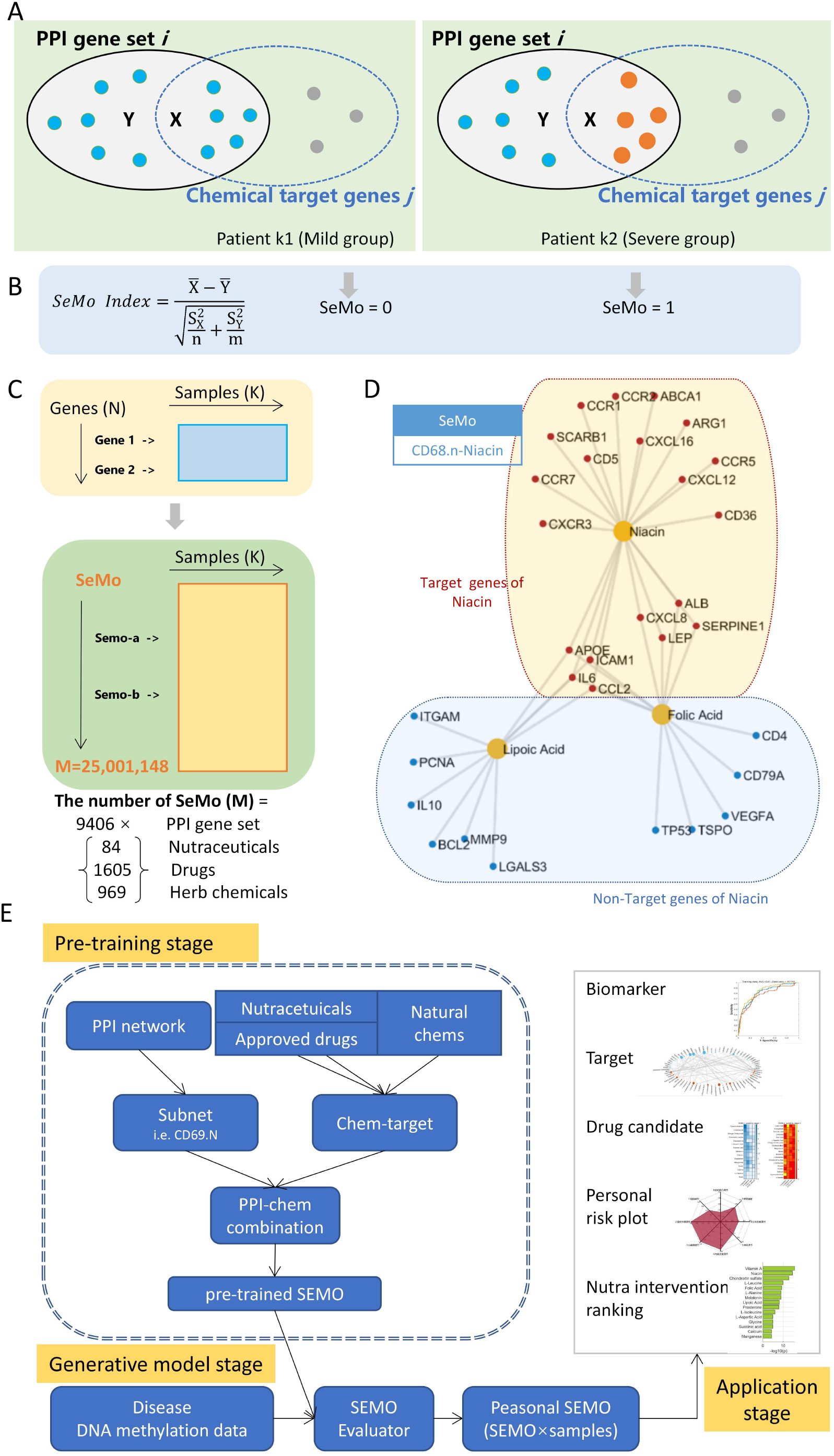
Pre-trained SEMO model generation and the whole information processing workflow. **A**. SEMO calculation is initiated by finding the intersection of the genes in the protein-protein interaction network and chemical target genes. **B**. The SEMO index is then calculated through the T-TEST between two sets – set x and set y. **C**. A SEMO matrix can be generated by applying this calculation to patient samples. **D**. An example of a SEMO: CD68.N-Niacin. **E**. The whole information processing workflow is divided into three stages: pre-training stage, generative model stage, and application stage.

In a whole-genome DNA methylation microarray data, we transformed the methylation sites × samples matrix into a genes × samples matrix by generating a representative methylation score for each gene. After iteratively combining each PPI gene set with chemical gene sets, we generated a SEMO× samples matrix (**Figure 1C**). In theory, there are 25,001,148 SEMO features that can be generated from 9406 PPI gene sets with ≥ 20 genes, combined with 2658 chemical gene sets. This allowed us to transform each patient’s DNA methylation profiling into a personal SEMO pattern. To reduce the computation cost, we filtered out a core set of phenotype associated PPI by calculating the association of PPI sets with COVID-19 severity (**Methods**).

To demonstrate the potential effects of targeted chemicals acting on a protein-protein interaction (PPI) network, **Figure 1D** presents an example of the SEMO generated from the CD68.N and niacin gene sets. The proteins depicted in the figure are all members of the CD68.N gene set, with CD68 being a marker molecule of macrophages. Genes surrounded by orange boxes indicate those that are the target of niacin, whereas those surrounded by blue boxes indicate those that are not the target of niacin. The SEMO index is calculated to determine the statistical difference between gene sets that are targets of the studied chemical and those that are not.

In order to screen and identify SEMOs related to phenotype, a complete information processing workflow and algorithms are illustrated in **Figure 1E**. The process includes three main phases: pre-training phase, generative model phase, and application phase. In the pre-training phase, the required combinations of PPIs and chemicals are generated. In the generative model phase, based on DNA methylation data, each individual’s gene-level data is transformed into SEMO dimensional data by SEMO calculation. The SEMO Evaluator is a critical data processing unit here, used to calculate the correlation between SEMO Index and phenotypes. Through this unit, SEMOs associated with phenotypes are formed and encoded into a SEMO Network -- a network that encodes relationships among PPIs, chemicals, and phenotype.

In the application phase, several potential outputs can be generated, such as biomarkers or predictive models for phenotype, as well as therapeutic or intervention targets identified from the most frequent appearing SEMOs associated with phenotype. Additionally, repurposing drugs or candidate drugs can be extracted from these same SEMOs, along with personal risk plots and personalized therapy or intervention recommendations (**Figure 1E**).

### Nutraceuticals impact on key immune genes networks during COVID-19

We examined whether the SEMO method could uncover the interaction between essential immune gene networks and nutraceuticals with respect to Covid-19. To do this, we initially studied the correlation of Covid-19 with a limited set of PPIs including CD68.N, a signature of macrophages and an important factor in long COVID; CD4.N, a signature of CD4+ T cells; IL6.N, associated with cytokine storm; CXCL8.N, correlated with both cytokine storm and disease severity; and FOXP3.N, the unique gene set for Treg cells.

**Figure 2A** shows the 4 most significant SEMO indices associated with disease severity in the CD68.N gene set. According to the 2-class t-test p-value of the SEMO indices in the severe group and mild group, the most significant SEMO indices were CD68.N-Lipoic Acid, CD68.N-Succinic Acid, CD68.N-Magnesium, and CD68.N-Niacin. An analysis of the odds ratios of single SEMO classifiers and patient groups (severe/mild) was also performed. For CD68.N-Lipoic Acid, the average SEMO index in the severe group was found to be significantly lower than that of the mild group (p=3.3e-14). Upon further examination of the gene set (the overlap between CD68 network and Lipoic Acid target genes), IL6 and IL10 were included. This indicates that Lipoic Acid could perturb the CD68.N gene set by affecting the DNA methylation levels of IL6 and IL10, two key markers in cytokine storms related to COVID-19. Lipoic Acid has also been proven to have beneficial effects on COVID-19 severity by inhibiting the overproduction of reactive oxygen species (ROS) and inflammatory cytokines [10, 11].

**Figure 2.**
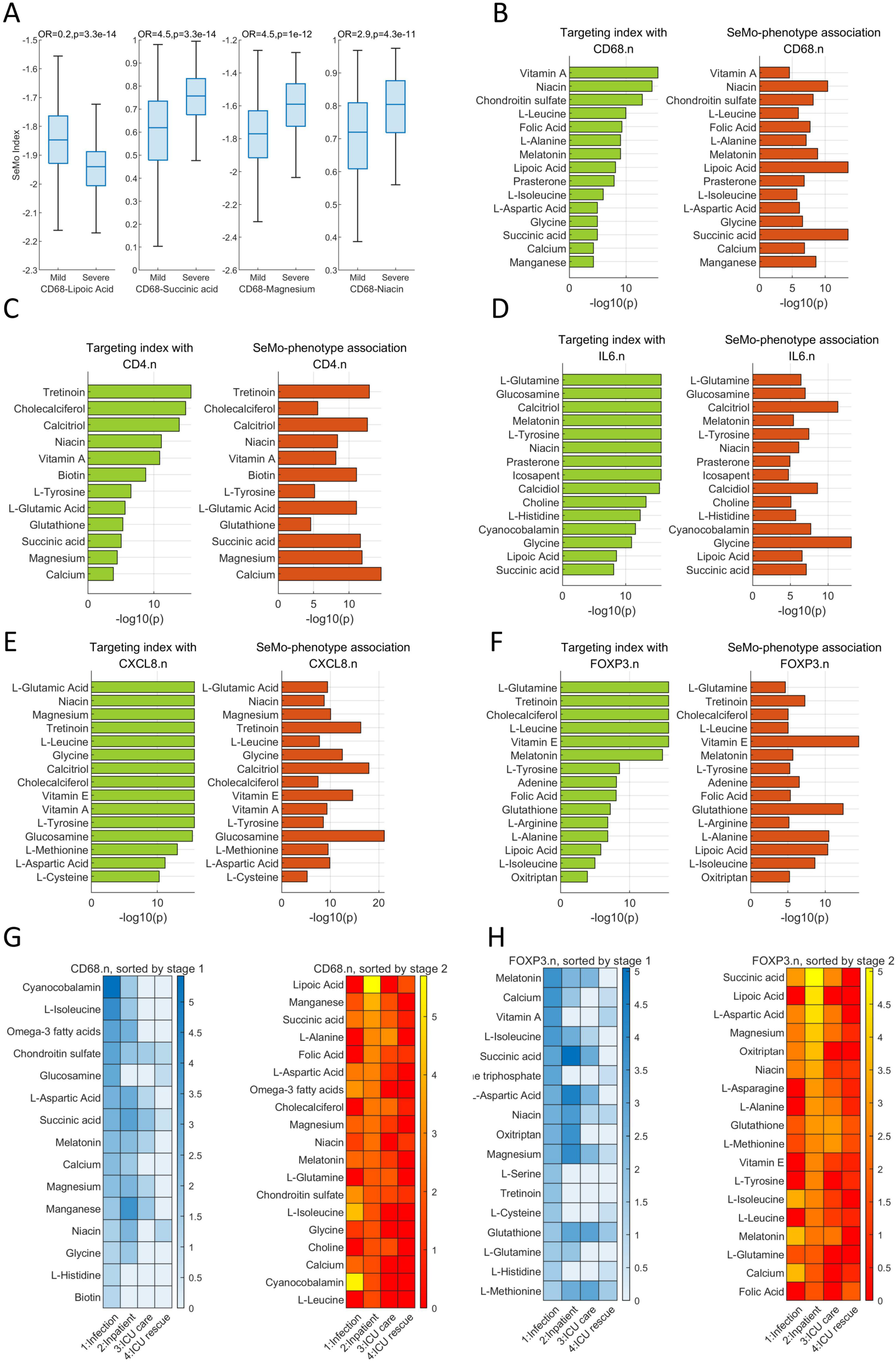
Key immune function related SEMO features associated with Covid-19 disease severity. **A**. The most statistically significant CD68 related SEMO features identified in severe versus mild groups in data 1. The combination of CD68 gene set with different nutraceuticals gene sets were calculated for SEMO features, and the differential significance of SEMO features in severe versus mild groups were assessed using two-class t tests with associated p values reported. **B**. The interaction of CD68 gene network with nutraceutical gene set is presented, with the left subplot displaying the gene set overlap significance (-log10(p), p is the hypergeometric test p value) between CD68 gene set and multiple nutraceuticals gene sets, and the right subplot showing the association of CD68-nutraceuticals SEMO features with severity. **C-F**. This presentation is also used to depict the interaction of CD4, IL6, CXCL8 and FOXP3 gene networks with nutraceuticals. **G**. The importance score of CD68-nutraceuticals SEMO in different stages of Covid-19 infection (positive, inpatient, ICU care and ICU rescue) is shown in a heatmap, with the left subplot sorted by the importance score in stage 1 and the right subplot sorted by the importance score in stage 2. **H**. The importance heatmap for FOXP3 gene network.

Compared to Lipoic acid-related SEMO, the other three top severity-associated nutraceuticals showed higher levels of SEMO index in the severe group than the mild group. This might suggest that these nutraceuticals have different mechanisms of action compared to Lipoic acid. Take Niacin as example, the niacin supplementation increase the nicotinamide adenine dinucleotide (NAD+) and operates as a metabolic switch to specify macrophage effector responses [12]. The SEMO calculation and phenotype association calculation present here provide a unique opportunity to generate testable hypothesis for chemical mechanism of action on immune systems.

**Figure 2B** (left subplot) displays the rank of gene set overlap significance (−log10(p), p being the hypergeometric test p-value, **methods**) between CD68 gene set and multiple nutraceuticals gene set. The most significant associated nutraceutical was vitamin A (1st line), followed by niacin, chondroitin sulfate, l-leucine and folic acid. This type of association is only related to the generic gene set of protein-protein interactions and nutraceutical target genes; thus, we refer to it as “targeting index”, which represents the baseline potential of PPIs that can be regulated by chemicals. Depending on the pathophysiological process, PPIs may be regulated by different sets of chemicals. The rank of SEMO-phenotype association (−log10(p), p being the 2-class t-test p-value in severe vs mild patients in data 1) is illustrated in **Figure 2B** (right subplot). In this case, the SEMO derived from the combination of vitamin A (the top 1 nutraceutical for CD68.n targeting index) with CD68.n showed a relatively small association with disease severity. The top significant SEMO was derived from lipoic acid, succinic acid and niacin. Succinate has also been reported to play an important role in regulating inflammation in macrophages [13].

Similar analyses were conducted on CD4.n, IL6.n, CXCL8.n and FOXP3.n. Briefly, the most severity-associated nutraceuticals for CD4.n were calcium, glycine for IL6.n, glucosamine for CXCL8.n and vitamin E for FOXP3.n (**Figure 2C-2F**).

The results presented here suggest that multi-stage SEMO analysis has the potential to generate useful hypothesis concerning the nutritional requirements of different patient groups. The nutraceutical-PPI pair may also play different roles in multiple stages of disease progression. To illustrate this, a multi-stage SEMO-phenotype heatmap was constructed for CD68.N (**Figure 2G**). The small squares in the heatmap represent the 2-class t-test p-value (-log10(p)) between the infection and negative patients (Stage 1), inpatient and home care (Stage 2), ICU and inpatient (Stage 3), and dead and ICU (Stage 4). When the lines (nutraceuticals) were ranked by important score in Stage 1 (left subplot), cyanocobalamin (Vitamin B12) was the most associated. Cyanocobalamin, l-isoleucine, and omega-3 were only important in Stages 1 and 2. However, some nutraceuticals were important across all four stages, such as chondroitin sulfate (Line 4) and succinic acid (Line 7). When the lines were sorted according to Stage 2 (right subplot), lipoic acid was ranked 1st. A similar analysis for Foxp3.N was conducted with melatonin as the most important nutraceutical for Stage 1 and succinic acid for Stage 2 (**Figure 2H**).

### Exploring protein-chemical-phenotype interactions using SEMO networks

By combining PPI gene sets with nutraceuticals, we scanned the associations between the resulting SEMO indices and severity in Data 1. After analyzing all SEMO features, we were able to identify the difference between severe and mild cases. If a PPI-chemical pair derived SEMO revealed a significant association with phenotype, we could construct a new type of network which links the given PPI to the corresponding chemical. The traditional PPI subnetworks (e.g. CD68.N) were represented here by a single node, thus forming a higher order network based on the PPI network (**Figure 3A**). In the SEMO network generated by the PPI-nutraceutical in Data 1, several central nodes, ITGAM.N, MMP9.N, and MMP2.N emerged, suggesting that these protein subnetworks are relevant to determining COVID-19 severity and thus could be targeted by multiple nutraceutical interventions.

**Figure 3.**
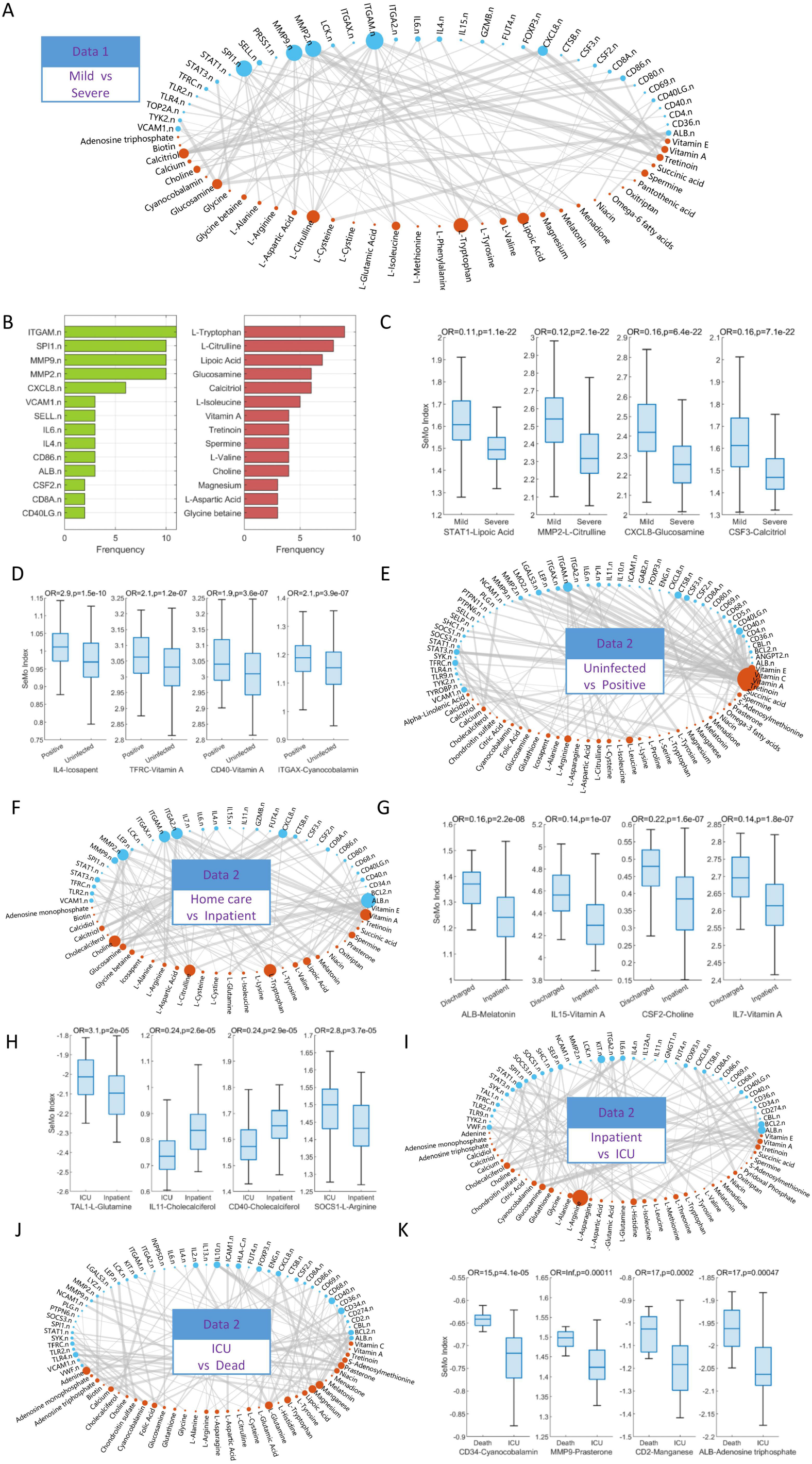
Exploring protein-chemical-phenotype interactions using SEMO networks. **A**. An inter-connecting relationship network between PPI gene sets and nutraceuticals from the top 100 Severity associated SEMO features, with node size reflecting the node degree, and connecting line thickness reflecting the chemical-PPI targeting significance (-log10(p)) as described in the Methods section. **B**. The top frequent PPI gene sets and nutraceutical that appear in the top 200 Severity associated SEMO features in Data 1. **C**. The four most significant SEMO features identified in the Severe vs Mild group of Data 1. **D-K**. The top associated SEMO features and SEMO networks connecting PPI sets to chemicals for various phenotypes, such as Uninfected vs Positive, Home care vs Inpatient, Inpatient vs ICU, and ICU vs Dead.

We ranked all SEMO features by the two-class t-test p-value (severe vs. mild group) and calculated the frequency of all PPIs and chemical sets appearing in the top 200 SEMO features in Data 1 (**Figure 3B**). The top 1 frequent PPI was the ITGAM gene network, in which the hub gene was ITGAM, the integrin subunit alpha m. ITGAM is important in the adherence of neutrophils and monocytes to stimulated endothelium and has been reported to be associated with long-term pulmonary complications in post-COVID-19 patients [14], suggesting that this gene network is important in COVID-19 severity. MMP9 (matrix metalloproteinase 9) and MMP2 networks, which are important in regulating inflammatory responses, have also been reported to be associated with mortality in COVID-19 patients [15]. Vascular cell adhesion molecule 1 (VCAM-1), which mediates leukocyte-endothelial cell adhesion, has been reported to be associated with unfavorable outcomes on the general ward and death on the ICU [16]. Another group of high frequent PPIs included inflammatory cytokines such as CXCL8, IL4, and IL6, and the dysregulation of these protein networks might also contribute to the inflammatory syndromes of the central nervous system in COVID-19 patients [17]. CD8A and GZMB gene networks, which are closely associated with CD8+ T cell functions and COVID-19 disease severity [18],were also highlighted. These PPIs suggest that T-cell activation, endothelium regulation, and cytokine/chemotaxin-related processes are important in COVID-19 severity and could be regulated by nutraceutical intervention. The importance of these biological processes is consistent with several hallmarks analysis of COVID-19 severity [3, 16].

The high frequent chemicals identified in the PPI-chemical networks of severity-associated metabolites (**Figure 3B**) revealed tryptophan as the top-ranked metabolite, which is in accordance with reports indicating its accumulation to be important for critically ill COVID-19 patients [4]. Additionally, citrulline was reported as an indicator for COVID-19 outcome [19] and l-isoleucine was significantly associated with disease severity and the gut microbiome of those affected by COVID-19, exhibiting impaired capacity for l-isoleucine biosynthesis [20].

While this list of metabolites serves as potential biomarkers for the disease, it also highlights a group of modulators for regulating the severity of or treating COVID-19. Several top-ranked nutraceuticals, such as lipoic acid [10, 11], N-acetyl-D-glucosamine [21], calcitriol (vitamin D3) [22], vitamin A/tretinoin [23], and spermine [24], have been suggested to have beneficial effects when used in symptom management and treatment of COVID-19.

For all combinations of PPIs and nutraceuticals, the top 4 most significant SEMO associated with disease severity in Data 1 were shown in **Figure 3C**. The SEMO calculation pointed out the potential target PPIs, i.e., STAT1 by lipoic acid, MMP2 by citrulline, CXCL8 by glucosamine, and CSF3 by calcitriol (vitamin D3). The SEMO index of STAT1-lipoic acid was significantly lower in the severe group compared to the mild group, with a two-class t-test p-value of 1.1e-22. Using the mean of this SEMO as a cutoff, the SEMO-high group (SEMO >= mean) had a lower risk of severity than the SEMO-low group, with a risk odds ratio of 0.11.

### Identifying metabolite biomarkers and modulators for COVID-19 severity

By comparing the SEMO features between different patient groups in data set 2, we identified statistically significant SEMO features associated with different stages of COVID infection. In the uninfected vs positive patient group (**Figure 3D**), the most connected chemical was Vitamin A (the biggest red node in the eastern direction of the network, **Figure 3E**), and TFRC-Vitamin A and CD40-Vitamin A were listed in the top 4 significant SEMO features in the positive vs uninfected patient group (**Figure 3D**). The unique importance of Vitamin A in the early stages of infection may be related to its unique role in the respiratory tract and in supporting the repair of the respiratory epithelium [23].

In the home care vs inpatient patient group, the important chemical sets were Vitamin A, L-Tryptophan and L-Citrulline (**Figure 3F**), and two Vitamin A-involved SEMO features, IL15-Vitamin A and IL7-Vitamin A, were listed in the top 4 significant SEMO features (**Figure 3G**). At this stage, ALB.N (Albumin) was the most highlighted PPI set (**Figure 3F**).

In the inpatient vs ICU patient group, the most important chemical was L-Arginine (**Figure 3I**), and SOCS1-L-Arginine was listed in the top 4 significant SEMO features (**Figure 3H**). Arginine is important as both a severity biomarker and a potential therapeutic option. First, arginine depletion, together with tryptophan metabolites accumulation, was reported to be correlated with T-cell dysfunction in critical COVID-19 patients [4]. Secondly, in terms of therapeutic implications, a potential regulatory crosstalk between arginine, tryptophan, purine metabolism and hyper inflammation in COVID-19 patients was reported, and targeting metabolism significantly modulates the pro-inflammatory cytokines release by peripheral blood mononuclear cells [25]. Thirdly, clinical studies showed that combining L-Arginine with other nutrients such as Vitamin C can improve long-COVID symptoms [26].

In the ICU vs dead patient group, the important chemical sets were Manganese, Magnesium and L-Glutamine (**Figure 3J, 3K**). Lower levels of manganese, magnesium and zinc were reported in severe patients compared to mild COVID-19 patients [27].

### Dynamic changes of PPI-Chemical interaction in COVID-19

To enable comparison of the dynamic changes of relative importance of the same protein networks, we generated an importance heatmap of various PPI gene sets. Based on Data 2, the frequencies of each PPI set (representing the relative importance) appearing in the top 200 significantly differential SEMO features for four stages (uninfected vs infection, home care vs inpatient, inpatient vs ICU, ICU vs death) were plotted as a heatmap (**Figure 4A**). Here, only the PPI sets where the sum of relative importance of stage 1: infection and stage 2: inpatient is larger than 5 were plotted, and the lines of PPI sets were sorted by the relative importance in stage 1: infection.

**Figure 4.**
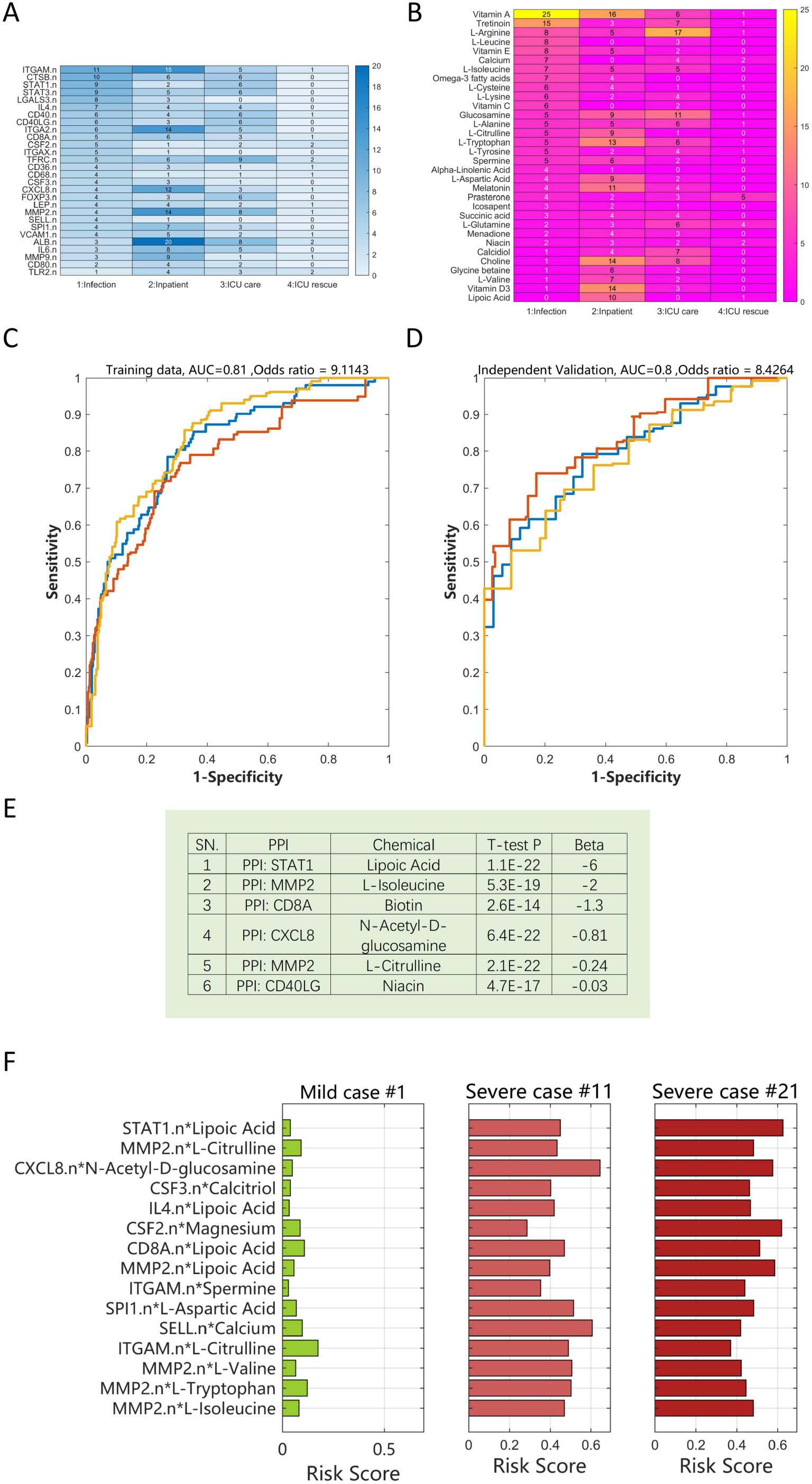
SEMO analysis for stage-specific nutraceutical demands, severity predictions, and personalized risk profiling. **A**. The importance score of PPIs in different stages of COVID-19 infection (positive, inpatient, ICU care, and ICU rescue). The heatmap was used to present the importance score of each drug, which is the frequency of it appearing in the top 200 phenotype-associated PPIs-drug SEMO (phenotype was defined as Stage 1: infection group vs negative group, Stage 2: inpatient vs home care, Stage 3: ICU vs inpatient, and Stage 4: dead vs ICU). The PPI list was sorted by their importance score in Stage 1. **B**. The importance score of nutraceuticals in different stages of COVID-19 infection was also generated. The nutraceutical list was sorted by their importance score in Stage 1. **C**. Severity prediction performances in training data (data 1) and independent validation data (data 2) were presented in the form of ROC curves and AUCs as well as odds ratios of prediction models. **D**. Severity prediction performance in independent validation data (data 2). ROC curve, AUC and odds ratio were shown. **E**. The model parameters of the best model identified from the Lasso machine learning algorithm. **F**. Personalized risk plots based on SEMO features were illustrated. The left subplot showed the risk profiling of a patient from the mild group, where the corresponding risk score could be calculated for each SEMO feature (see **Methods**). The center and right subplots illustrated risk plots of two patients from the severe group.

Firstly, we compared which SEMO of Data 2 is similar to the severe vs mild stage of Data 1. Previously, we discussed the significantly differential SEMO features in the severe vs mild patient group in Data 1, the most important PPI sets being ITGAM.N, SPI1.N, MMP9.N, MMP2.N and CXCL8.N. In this four stage plot (**Figure 4A**), the column of stage 2: inpatient is the most similar top important list of PPI sets, comprising ITGAM.N (importance score 15), MMP2.N (14), CXCL8 (14), MMP9.N (9) and SPI1.N (7). This suggests that the stage 2: inpatient (inpatient vs home care) in Data 2 is relatively similar to the severe vs mild comparison in Data 1.

Secondly, we examined the top list of PPI sets in stage 1: infection. ITGAM.N ranked 1st in stage 1 (importance score = 11) but was even more important at stage 2: inpatient (importance score = 15). The 2nd line of the heatmap, CTSB.N, showed a pattern where it is most important at stage 1 (**Figure 4A**). Interestingly, CTSB (cathepsin B) and CTSL (cathepsin L, one of the key members of CTSB.N protein network) were reported to facilitate the entry of SARS-CoV-2 into the human host cell by cleaving the spike protein of the virus [28, 29] and are potential therapeutic targets for Covid-19 [29, 30].

The fifth line, LGALS3.n, represents a protein network that was important in both stage 1 and stage 2 (**Figure 4A**). LGALS3, also known as galectin-3, was recently reported in a Phase IB/IIA randomized controlled clinical trial as a potential target in COVID-19 pneumonitis [31]. CD40.n and CD40LG are both important in stages 1 to 3 (the seventh and eighth line), suggesting that this protein interaction is relevant to COVID-19 pathophysiology. Studies have demonstrated that the CD40L/CD40 interaction is involved in platelet-related inflammation processes [32]. In severe COVID-19 patients, hyper coagulation, monocyte activation, and cytokine storms are correlated, and the CD40L/CD40-related process is considered a potential therapeutic target [33].

In short, the heatmap in **Figure 4A** suggests that SEMO calculation could reveal important pathophysiological processes and key therapeutic targets, at least in immune cell and endothelium-related processes.

For chemicals, we also illustrate the importance profiling across four stages. As previously discussed, vitamin A/tretinoin are the most important chemicals in stage 1 and L-arginine is highlighted in stage 3 (**Figure 4B**).

### SEMO based COVID 19 disease severity prediction model

Based on the predictive capacity of SEMO features, we constructed multiple parameter prediction models using a machine learning algorithm (LASSO) to classify patients as either severe or mild groups. The AUC in the training data was 81% (with an odds ratio of 9.11, **Figure 4C**) and 80% in the independent validation set (with an odds ratio of 8.43, **Figure 4D**). This relatively stable AUC suggests that our model is relatively robust. The optimal model used six SEMO features, STAT1.n-Lipoic acid, MMP2.n-L-isoleucine, CD8A.n-Biotin, CXCL8.n-Glucosamine, MMP2.n-L-Citrulline, and CD40LG.n-Niacin (**Figure 4E**).

### SEMO based personal risk profiling and nutraceutical regimes

In order to establish personalized nutrition interventions for the prevention of severity, we have developed a “personal risk profiling” tool (see **Methods**). This tool can illustrate the relative severity risk from various SEMO features (**Figure 4F**). In the left panel, the horizontal bar graph shows the relative risk of severity stemming from a patient in the mild group, with its riskiest bar showing a value of less than 0.1. The center and right panels of **Figure 4F** illustrate the risk profiles of two patients in the severe group, with the riskiest bar being greater than 0.4. Even among people in the same severe group, there may be slight differences in terms of the most important risk factors. In the center panel, the top risk is from the feature “CXCL8.n-glucosamine” while in the right panel, the two most important risk factors are “STAT1.n-lipoic acid” and “CSF2.n-magnesium”. With this information, personalized nutraceutical regimes can be generated tailored to each individual, such as the combination of lipoic acid and magnesium for the patient shown in the right panel.

### SEMO index are associated with immune cells composition

The SEMO features associated with immune cells composition were further evaluated. An immune profiling data set, which included both the immune cell proportions and genome-wide DNA methylation profiling data, was used to calculate the Pearson correlation coefficient (ρ) and p-value between each SEMO feature and 12 cell type proportions (**Methods**). These cell types included neutrophils, eosinophils, basophils, monocytes, B cells (naïve and memory), CD4+ and CD8+ (naïve and memory) cells, natural killers, and T regulatory cells. Additionally, some derived parameters such as the neutrophils to lymphocytes ratio (NLR), the monocytes to lymphocytes ratio (MLR), and the CD4/CD8 ratio were also calculated.

The correlations between the severity associated SEMO features and the 12 cell types were evaluated (**Figure 5A**). For each SEMO, only the cell type with the highest correlation coefficient was shown. Most of the severity associated SEMO features were found to be correlated with CD4+ T cell proportions; for example, STAT1-lipoic acid and CSF3-calcitriol (vitamin D3) suggest potential beneficial effects of these nutraceuticals related to CD4+ T cells. Additionally, some SEMO features were associated with monocytes, such as MMP2-L-citrulline and CXCL8-glucosamine.

**Figure 5.**
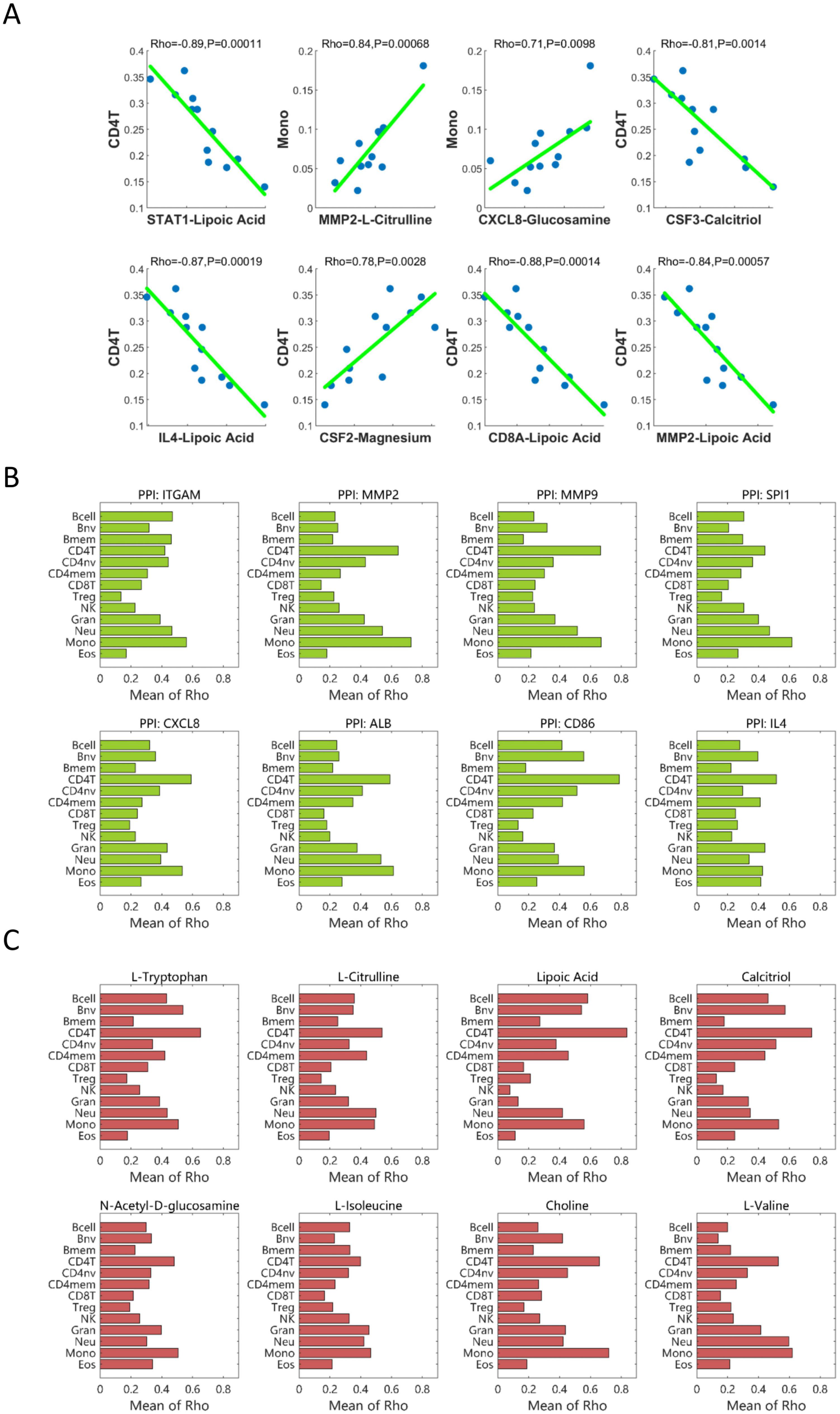
The association between severity associated SEMO features and immune cell proportions. **A**. The linear correlations between each SEMO feature (x-axis) and its respective immune cell proportion (y-axis). **B**. The correlation of PPI gene set with different immune cell proportions. Taking “ITGAM” as an example. We extracted all SEMO features containing “ITGAM”, and then calculated the mean Pearson Correlation Rho values between the SEMO features and each cell type. **C**. The correlation of chemicals with different immune cell proportions. Using “L-Tryptophan” as an example, we collected all SEMO features containing “L-Tryptophan” and computed the mean Pearson Correlation Rho values between these SEMO features and each cell proportions.

To further investigate which type of immune cells were correlated with specific PPI gene sets, for example ITGAM.N, all severity associated SEMO features that included this PPI (ITGAM.N-spermine, ITGAM.N-L-citrulline, ITGAM.N-L-tryptophan, etc.) were queried and the mean value of the Pearson correlation coefficient between these SEMO features and each cell proportion was calculated (**Figure 5B**). For ITGAM.N, monocytes were the most correlated cell type. For MMP2.N and MMP9.N, monocytes were the most correlated cell types, with CD4+ T cells ranked second. For CD86.N, CD4+ T cells were the most associated cell type.

For each chemical, the correlated cell types were investigated by similar calculations (**Figure 5C**). For the top 8 severity associated chemicals, the most two correlated cell types were CD4+ T cells and monocytes, suggesting that the potential modulation of these chemicals was mostly associated with changes in the cell count of these two types of cells. Additionally, other cell types were shown with relatively high correlations; for example, naïve B cells were the second correlated cell type for L-tryptophan and calcitriol (vitamin D3), whereas neutrophils were the second correlated cell type for L-citrulline and granulocytes for L-isoleucine.

### Drug re-purposing opportunities for various stages of COVID 19

We applied SEMO analysis on FDA-approved drugs, and the top four significant severity-associated SEMO features were related to the drug heparin (**Figure 6A**). Furthermore, the targeting PPIs set for this analysis were MMP9 and CXCL8. Heparin was reported to affect matrix metalloproteinase and could potentially benefit severe COVID-19 patients through anticoagulation, as suggested by clinical trials[34, 35]. Additionally, SEMO MMP2.n-indomethacin were significantly differentiated in severe versus mild patients, with a p value of 1.2e-20 (**Figure 6A**). Indomethacin, an anti-inflammatory drug and broad-spectrum antiviral agent, has been reported to be safe and effective in treating COVID-19 patients in a randomized clinical trial [36]. In the list of the top 200 significant severity-associated SEMO by PPIs-drugs combination, the most frequent PPIs set was similar to SEMO by PPIs-nutraceuticals, and MMP9, MMP2, CXCL8, CD8A, among others, were also present in the list (**Figure 6B**, left panel).

**Figure 6.**
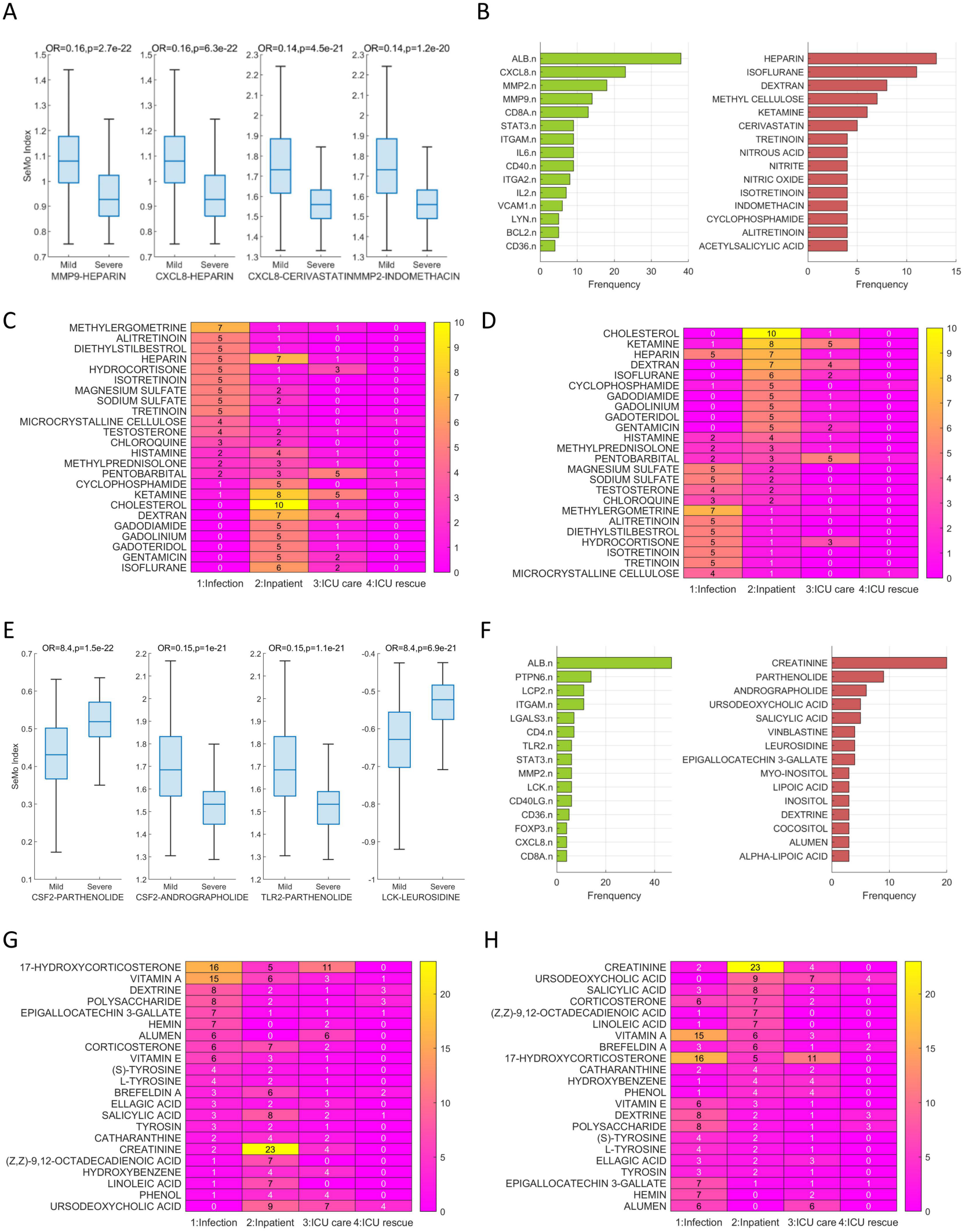
SEMO analysis for drug repurposing and discovery of functional chemicals from natural sources for COVID-19. **A**. The top four PPI-drug SEMO features that exhibited significant differences between the Severity and Mild groups in data 1. **B**. The most frequent PPI gene sets and drugs that appeared in the top 200 Severity-associated SEMO features in data 1. **C**. The drug importance scores across different stages of COVID-19 infection (positive, inpatient, ICU care, and ICU rescue). The drugs were sorted by their importance score in Stage 1. **D**. Another version of Subplot C with the drugs sorted by their importance score in Stage 2. **E**. The top four natural chemicals derived SEMO features that exhibited significant differences between the Severity and Mild groups in data 1. **F**. The most frequent PPI gene sets and natural product chemicals present in the top 200 Severity-associated SEMO features in data 1. **G**. The importance score of natural product chemicals across different stages of COVID-19 infection. The chemicals were sorted by their importance score in Stage 1 **H**. Another version of subplot **F**, with the chemicals sorted by their importance score in Stage 2.

Based on the patient group in data 2, we checked the importance scores of drugs in different stages of COVID-19. Here, the importance score is the frequency of drugs that appear in the top 200 phenotype-associated SEMO. The top drugs sorted by importance score in stage 1 are shown in **Figure 6C**. The drugs in this list indicate the mechanisms related to the first-line response to viral infection. For example, tretinoin (**Figure 6C**, right panel, line 2), also known as all-trans retinoic acid, is a natural metabolite of vitamin A. It was reported to inhibit SARS-CoV-2 infection by interrupting spike protein-mediated cellular entry [37]. In this case, alitretinoin, isotretinoin, and tretinoin were listed in lines 2, 6, and 9 (**Figure 6C**).

The top drugs sorted by importance score in stage 2 are shown in **Figure 6D**. Similar to the results in data 1, heparin scored high and was ranked 2 in the stage 2 column.

### Discovery of functional chemicals from natural sources for COVID-19

Nature products, herbs used in Traditional Chinese Medicine (TCM), have the potential to act as drug candidates against various stages of COVID 19. In particular, CSF2-Parthenolide, CSF2-Andrographolide, TLR2-Parthenolide and LCK-Leurosidine have been identified as the most significant severity associated SEMO features (**Figure 6E**). Parthenolide, which is the main active ingredient of the herb feverfew (Tanacetum parthenium), is known to have anti-inflammatory properties by inhibiting IL-6 production, as well as inhibitory activity on SARS-CoV-2 papain-like protease[38]. Andrographolide is the predominant active compound of the plant Andrographis paniculata, which is widely used in TCM and has anti-inflammatory functioning, with the potential to reduce cytokine storms[39]. Thus, these chemicals may affect virus responses by regulating CSF2 and TLR2 protein networks to regulate cytokine production.

On ranking the high frequent chemicals in data 1 within the top 200 severity associated SEMO groups (**Figure 6F**, right panel), Parthenolide is found at the second position, with Andrographolide at third place. Ursodeoxycholic acid is a naturally occurring bile acid that recently has been reported to be effective in reducing SARS-CoV-2 infection due to its regulation of the ACE2 pathway [40]. In the present results, SEMO CD4.n-Ursodeoxycholic acid and CD69.n-Ursodeoxycholic acid were observed to have a significantly differentiating effect between severe and mild cases, with 2-class t-test p-values of 1.1E-17 and 4.3E-17 respectively. This result suggests that Ursodeoxycholic acid might target CD4+ and CD69+ cells.

Various drugs may have varying levels of importance in different stages of COVID 19. Taking data 2 as an example, Epigallocatechin 3-gallate (EGCG) extracted from green tea was the fifth ranked drug in stage 1 (**Figure 6G**), whereas Ursodeoxycholic acid was ranked second in stage 2 (**Figure 6H**).

## Discussions

This paper presents a novel generative model to identify biomarkers and drug candidates as well as personalized therapeutic regimes in COVID-19. The approach hinges on the major steps of pre-training phase, generative model phase and application phase, involving the integration of data from protein-protein interactions (PPIs) and chemicals to transform individual gene-level data into SEMO-dimensional data by SEMO calculation.

The advantages of this method is that the SEMO network, as a generated model, could serve as an infrastructure to offers the opportunity to generate a wide range of outputs, including both biomarkers and therapeutic targets, as well as candidate drugs and personalized therapeutic regimes. This approach can identify not only gene modules with significant causal relationships as the disease progresses, but also compounds that may be able to regulate these specific signal modules. In addition, the application of this method provides insight into the dysregulation of immune signaling pathways in various COVID-19 stages, as well as the possible nutraceutical demands and modulatory effects of chemicals on immune cell signaling transduction activity. Moreover, the SEMO network’s higher-order knowledge representation of interactions between proteins, chemicals, and phenotype can help understand the association of SEMO with CD4+ T cells and monocytes proportions. Altogether, the results of this study strongly suggest that the generative model and SEMO network provide testable hypotheses for better management and treatment of long-term COVID-19 patient care, and open the door for further exploration and research.

In the field of computational biology, scientists have traditionally considered the biochemical pathways triggered by the binding of membrane receptors and ligands when studying disease and drug mechanisms. However, it can be challenging to model the effects of multiple drugs, targets, and their associated phenotypes based on these biochemical events, as comprehensive data collections are required. Here, we present the SEMO method which uses protein network modules as signal enhancers to create cumulative, complementary or synergistic regulation effects, making them biologically meaningful. This method is information-based and not necessarily chemistry-based.

Moreover, predicting disease outcome in elderly patients and the coexistence of multiple diseases remains a challenge as traditional models are usually developed for single diseases only. To address this, the World Health Organization (WHO) proposed the intrinsic capacity (IC) to quantify the capability of the human body. IC is more related to common mechanisms of multiple pathophysiology processes, and may therefore predict disease outcomes. However, traditional IC quantifications are often both time consuming and subjective due to questionnaires and professional assessments. We propose that a new type of intrinsic capacity quantification, which we call “capomics”, can be achieved by combining pre-trained models with phenotype specific omics data and generating a huge feature space to characterize the potential of an individual’s response to various disease-related factors. This is a data-driven method that only requires omics data as an input and does not require phenomics measurements. Capomics has a high dimensionality, providing a large set of generated features to characterize human capacity. For example, SEMO consists of a combination of PPI sets and chemical sets, which can help us identify phenotype-specific PPI and chemical modulators. The SEMO index of key immune-related protein networks such as CD68 and IL6 reflect the host’s ability to maintain proper balance in the immune process, either severely boosting or restraining inflammation. This SEMO framework could potentially serve as a fundamental map of intrinsic capacity, leading to greater insight into the development and progression of numerous diseases.

Nutritional factors have a significant impact on the immune system, making it a challenge to study multiple nutrients at once. This has limited the progress of research into nutrition’s role in Covid-19 management due to the complex relationship between multiple nutrients and the human immune system. This paper proposes a method to quickly generate hypotheses on the effects of multiple nutrients on immunity. DNA methylation profiling is seen as a powerful omics tool to reflect underlying mechanisms for human aging, immune remodeling, and environmental exposure, meaning that its rapid and cost-effective implementation could hold potential for patient management and clinical nursing.

The limitation of this paper is that the generative model is currently limited to the study of COVID-19, and further studies are needed to determine whether the model can be applied to other diseases. Additionally, the generative model is currently based on DNA methylation data, and its applicability to different types of data, such as gene expression data, needs to be explored in future studies. Additionally, the output results are reliant on the database used, particularly the data of PPI and protein-compound interactions, so further research into the impact of different databases on the results is necessary.

## Conclusions

In conclusion, this paper presents a novel generative model for identifying biomarkers, drug candidates, and personalized therapeutic regimes in COVID-19. The approach has the potential to generate a comprehensive range of outputs, understand dysregulation of immune signaling pathways, and inform possible nutraceutical demands and modulatory effects of chemicals on cellular signaling activity.

